# Exploring the role of competition induced by non-vaccine serotypes for herd protection

**DOI:** 10.1101/092593

**Authors:** G. L. Masala, M. Lipsitch, C. Bottomley, S. Flasche

**Affiliations:** London School of Hygiene and Tropical Medicine, London, UK; Harvard University, Boston, US

**Author notes:** Corresponding author: Stefan Flasche, London School of Hygiene and Tropical Medicine, WC1E 7HT London, UK.

**Keywords:** Pneumococcus, vaccination, serotype competition, herd protection

## Abstract

The competitive pressure from non-vaccine serotypes may have helped pneumococcal conjugate vaccines (PCVs) to limit vaccine serotype (VT) prevalence. We aim to investigate if, consequently, the indirect protection of higher valency vaccines could fall short of the profound effects of current formulations.

We compare three previously described pneumococcal models harmonized to simulate 20 serotypes with a combined pre-vaccination prevalence in <5y old children of 40%. We simulate vaccines of increasing valency by adding serotypes in order of their competitiveness and explore their ability to reduce VT carriage by 95% within 10 years after introduction.

All models predict that additional valency will reduce indirect vaccine effects and hence the overall vaccine impact on carriage both in children and adults. Consequently, the minimal effective coverage (efficacy against carriage * vaccine coverage) needed to eliminate vaccine type carriage increases with increasing valency. One model predicts this effect to be modest while the other two predict that high-valency vaccines may struggle to eliminate VT pneumococci unless vaccine efficacy against carriage can be substantially improved. Similar results were obtained when settings of higher transmission intensity and different PCV formulations were explored.

Failure to eliminate carriage as a result of increased valency could lead to overall decreased impact of vaccination if the disease burden caused by the added serotypes is low. Hence a comparison of vaccine formulations of varying valency, and pan-valent formulations in particular, should consider the invasiveness of targeted serotypes, as well as efficacy against carriage.

## Background

In 2000 the first pneumococcal conjugate vaccine (PCV), which provided protection against seven of the most pathogenic pneumococcal capsular serotypes, was licensed and recommended for immunization of infants in the US [1]. Subsequently, 10- and 13-valent formulations have been licensed and are now being used to prevent pneumococcal disease in more than 130 countries worldwide [2–11]. The incidence of carriage and disease associated with vaccine type serotypes (VT) declined in vaccinated children, and also in unvaccinated children and adults, after PCVs were introduced into national immunization programmes [12]. However, the overall prevalence of pneumococcal carriage remained approximately constant as non-vaccine serotypes (NVT), i.e., serotypes not targeted by the vaccine, filled the ecological niche [13]. The increased disease from these serotypes has partially offset the benefit of pneumococcal vaccination. As a result pneumococcal vaccines that target more or all serotypes are being developed [14].

Previous work has suggested that the competition between vaccine type (VT) and non-vaccine type (NVT) serotypes plays an important role in the herd protection observed in the post PCV era [15–17], In particular, by reducing acquisition of VT carriage, pneumococcal conjugate vaccines give NVTs a competitive advantage over VTs in the nasopharynx. Thus, in vaccinated populations the presence of NVT in vaccinated hosts provides additional competitive pressure, which combines with the immune protection afforded by the vaccine, to suppress VT colonization. Moreover, at the population level, there is competition between VT and NVT in non-vaccinated hosts, and the spread of VT is likely inhibited by competition from NVT in non-vaccinated people as well. In each of these cases, the mechanisms of competition might include direct competition in the nasopharynx [18], induction of innate immune effectors by NVT that also inhibit VT [19], and induction of forms of acquired immunity that also inhibit VT, such as Th17-based and antibody-based immunity to conserved antigens [20–22], For higher valent pneumococcal vaccines, including those that target proteins common to most pneumococci [14], this hypothesis implies that, by effectively losing the benefit of NVT competition, even with similar vaccine efficacy against pneumococcal carriage and disease, levels of indirect protection in unvaccinated individuals could fall short of the profound effects that have been observed with the routine use of conjugate vaccines.

In this paper we explored three previously developed dynamic modelling approaches for pneumococcal ecology as to whether they predict a similar contribution of NVT competition to the indirect effects of pneumococcal vaccination. We harmonized key model parameters that govern vaccine efficacy and pneumococcal epidemiology in the absence of vaccination and explored various vaccine scenarios to better understand the role of competition in providing protection.

## Methods

### Models

The model by Bottomley et al [16] (M_B) is a deterministic model that represents the pre-PCV steady state of pneumococcal infections in the Gambia and was used to predict the impact of introduction of PCV13 into the childhood vaccination program. It is fitted to local longitudinal pre-vaccination carriage data. Serotypes are grouped into three classes of transmissibility (low, medium, high) and clearance rate (high, medium, low).

Serotypes within the same group share the same properties. Pneumococcal carriers are assumed to gain partial immunity against acquisition of new serotypes during the episode of carriage, which represents the mechanism of serotype competition, and a proportion of clearances leads to life-long immunity against the cleared serotype, which balances competitive advantages to sustain serotype diversity. The default model parameterization is the same as reported in the original manuscript.

The model by Cobey & Lipsitch [23] (M_CL) is an individual-based model that represents a generic high-income setting. Serotypes differed by their intrinsic duration of carriage and their in vivo competitive ability. Pneumococcal carriers are assumed to gain partial protection, quantified by the competitive ability of the resident type, against acquisition of additional serotypes during the episode of carriage, which represents the mechanism of competition. Following clearance, the host’s susceptibility to any subsequent homologous acquisition as well as the duration of any subsequent carriage episode is reduced. The serotype-specific immunity accentuates within-serotype competition thus providing balancing selection on serotypes, and acquired immunity independent of capsule reduces fitness differences. The simulations presented in this work rely on the default model parameterization, i.e. homogeneous mixing, the default rate of acquisition of capsular immunity *σ* = 0.3) and the default rate of acquisition of nonspecific immunity assuming non-linear reduction in carriage duration (*∊* = 0.25).

The model by Flasche et al [15] (M_F) is an individual based model that generalizes the most commonly used deterministic pneumococcal model [17,24,25]. It represents a generic high income country setting. Serotypes differ by their intrinsic duration of carriage. Duration of carriage and susceptibility to acquisition decline with age but are exposure independent. The acquisition of a pneumococcus triggers both a transient homologous immune response, which represents serotype competition, and a transient heterologous immune response, which represents the mechanism to ensure serotype coexistence. Both immune responses are assumed to prevent additional acquisition of respective serotypes. Unless mentioned otherwise the simulations presented in this work rely on the default model parameterization, i.e. the duration of specific and non-specific immunity was 9 and 18 weeks respectively.

While there was no formal model selection process we included models that span most of the range of alternative dynamic pneumococcal model assumptions on serotype competition and natural immunity. An overview of the different modelling approaches in this study is shown in Table 1. The main differences between models in regards to this work are the assumptions on acquired immunity and the resulting mechanisms for competition and serotype coexistence.

**Table 1:** Summary of the main features of each model of pneumococcal transmission.

|  | Bottomley et al | Cobey & Lipsitch | Flasche et al |
| --- | --- | --- | --- |
| **Model type** | Compartmental | Individual based | Individual based |
| **Demographics** | Gambia | High income country | High income country |
| **Mixing patterns** | Homogeneous | Homogeneous | Age-assortative |
| **Natural immunity as a result of infection** | *non-specific*: transient immunity for the duration of infection<br>*anticapsular*: chance to develop permanent homologous immunity | *non-specific*: permanent increase in clearance rate, transient reduction in acquisition rate for the duration of infection<br>*anticapsular*: permanent reduction in susceptibility to homologous infection | *non-specific*: transient immunity to heterologous infection<br>*anticapsular*: additional transient immunity to homologous infection<br>*other*: exposure independent reduction of susceptibility and carriage duration with age |
| **Vaccine induced immunity** | Like anticapsular natural immunity but higher chance for protection | Like anticapsular natural immunity but stronger protection | Like anticapsular natural immunity but longer protection |

### Analyses

We harmonized models to simulate 20 artificial serotypes with a combined pre-vaccination prevalence in <5y old children of 40%, approximating a high income setting with moderate transmission, or 70%, approximating a low income setting with high transmission intensity. In M_B serotypes were evenly distributed between the three classes, i.e. 7, 7,6 serotypes of low, mid and high transmission intensity. For each model, parameters governing transmission intensity were scaled to achieve the desired prevalence. In M_F changing the transmission intensity alone was insufficient to achieve the targeted 70% prevalence (compare discussion in Flasche et al [15]). Hence, once the effects of increasing the transmission intensity saturated it was kept constant and the duration of specific and non-specific immunity were subsequently decreased to 6 and 12 weeks respectively to achieve the targeted prevalence. As M_B was not age structured we split all compartments into an <5y old and a 5 years and older compartment retaining all original parameters and constant rate of transition between the strata such that the mean time spent in the <5 stratum was 5 years. To minimize the effects of stochasticity M_CL was run 10 times and mean values are presented. Stochastic variability on presented outcomes was too small to visualize in the Figures so it was omitted. The Simpson Index was calculated as a measure of serotype diversity [26].

We explored two vaccines with alternative compositions each. Firstly, we compared generic vaccines of increasing valency where serotypes are included in order of their competitiveness. Serotype “1” is the strongest competitor, indicated in all models by the highest prevalence in the pre vaccine era, and “20” the lowest. Note that the serotype names do not correspond to the numbering conventionally used for pneumococcal serotypes. Secondly, to approximate PCV7, PCV10, PCV13 and PCV15 we modelled inclusion of serotypes into the vaccine in respect to their observed paediatric prevalence rank among carriage globally. For example PCV7 targets serotypes 4,6B,9V,14,18C,19F and 23F which are the 18^th^, 2^nd^, 6^th^, 5^th^, 11^th^, 1^st^, 3^rd^ most prevalent serotypes globally; hence the PCV7-like vaccine in this work targets model serotypes 1,2,3,5,6,11 and 18 (Table 2) [27], Serotypes of a lower rank than 20 were not included since the models only consisted of 20 serotypes.

**Table 2:** PCV formulations and the ranks of each serotype in terms of its global prevalence according to a review on the global distribution of paediatric pneumococcal carriage.

|  | Serotypes (rank) |
| --- | --- |
| **PCV7** | 4 (18), 6B (2), 9V (6), 14 (5), 18C (11), 19F (1), 23F (3) |
| **PCV10** | + 1 (31), 5 (38), 7F (32) |
| **PCV13** | + 3 (9), 6A (4), 19A (7) |
| **PCV15** | + 22F (27), 33F (24) |

M_B and M_F assume that both immunity, including vaccine induced immunity act as all-or-nothing while the Cobey and Lipsitch model assumes it is leaky. The models were run to predict 1) the impact of vaccination against each targeted serotype [28] (assuming 100% coverage and 55% efficacy) and 2) the effective coverage needed to achieve elimination of VT carriage. We defined effective coverage as vaccine efficacy times vaccine coverage (N.B. in the two models that assume all-or nothing vaccine protection this is equivalent to the fraction of the population that is protected by vaccination), and elimination of VT carriage as a reduction of VT carriers of 95% or more. The impact of vaccination is measured as either the percentage reduction in the number of VT carriers, or alternatively with any serotype, in year 10 after the start of vaccination if compared to the year before vaccination (steady state).

## Results

Models differed in the proportion of children among the simulated population. M_B assumed an age distribution based on the Gambia and hence that children younger than 5 years old make up 20% of the total population. In both other models that represent high income countries the corresponding proportion was 5% (Figure 1). Pneumococcal carriage prevalence was almost evenly distributed across serotypes in M_F and dominated by fewer serotypes in M_CL. The Simpsons index in children was 0.871, 0.845, 0.885 in the moderate transmission scenario for M_B, M_CL and M_F and 0.925, 0.862, 0.927 in the high transmission scenario. The models differed how the targeted prevalence in children translated into prevalence in older individuals (Figure 1). Carriage prevalence decreases with age except for M_B which was originally designed to be age independent and hence uses the same parameters for both age groups.

**Fig. 1.**
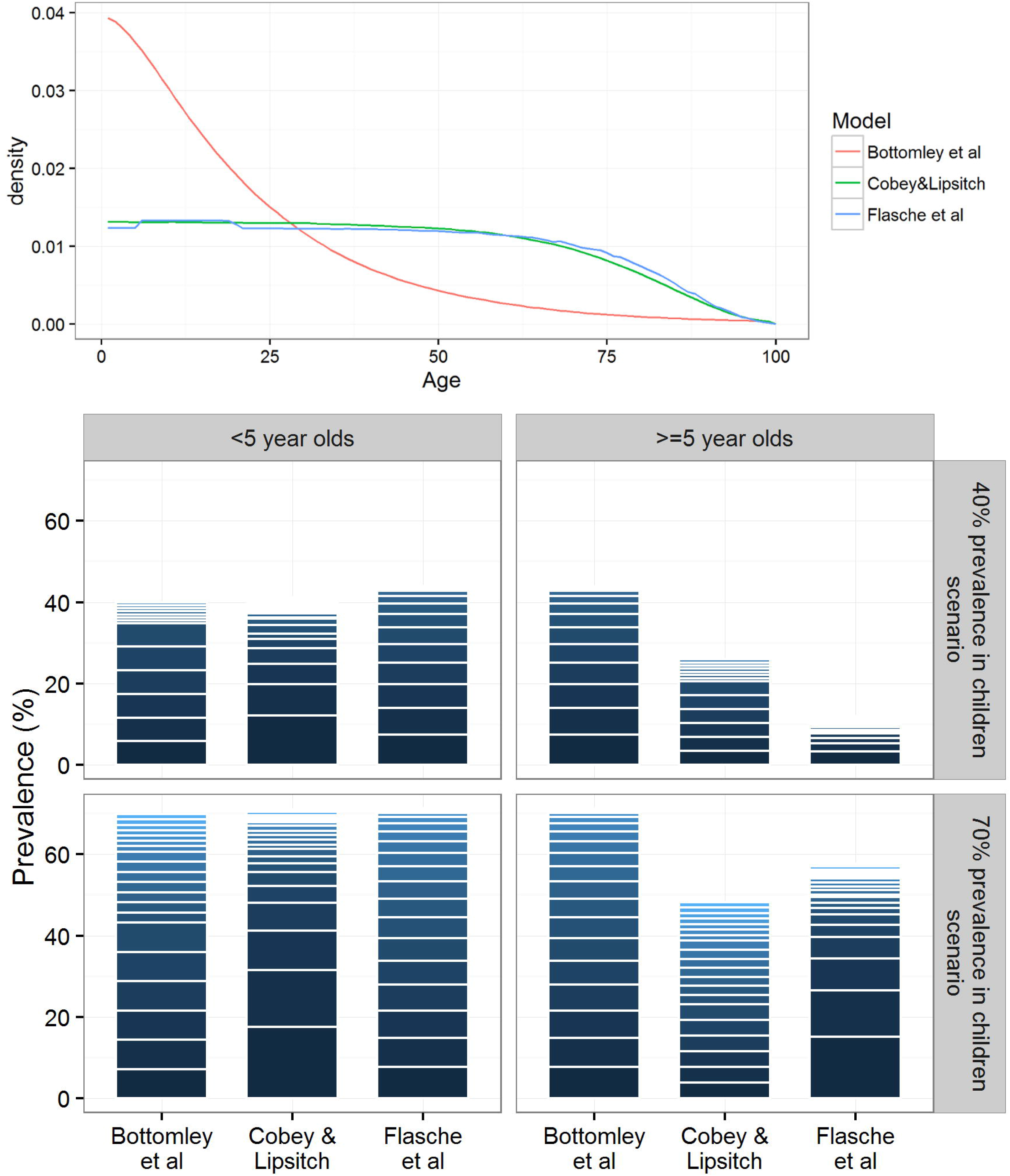
Model demographics and serotypes distribution before the introduction of vaccination. Upper panel: The cumulative age distribution of the model populations. Lower panel: a stacked barplot to illustrate the predicted serotype distributions (stacked prevalence of serotype specific carriage episodes scaled to the overall carriage prevalence) in children and the rest of the population in low and high transmission settings.

In the moderate transmission scenario, the vaccine impact against vaccine type carriage 10 years after the start of vaccination of a vaccine with 55% effective coverage decreased with increasing vaccine valency in all three models (Figure 2). The effect of including more VTs was least visible in M_CL where only inclusion of almost all serotypes (15 or more) reduced the impact on VT carriage to allow VT circulation. The M_B predicted the steepest decrease in vaccine impact on VT prevalence as a result of inclusion of highly and moderately competitive serotypes, however, further inclusion of weakly competitive serotypes which were hardly carried in this scenario did not change the impact of vaccination. Similar dynamics were observed for older individuals and the high transmission scenario, however, M_F predicted a small initial increase in vaccine impact on VT carriage before the impact decreased for higher valencies.

**Fig. 2.**
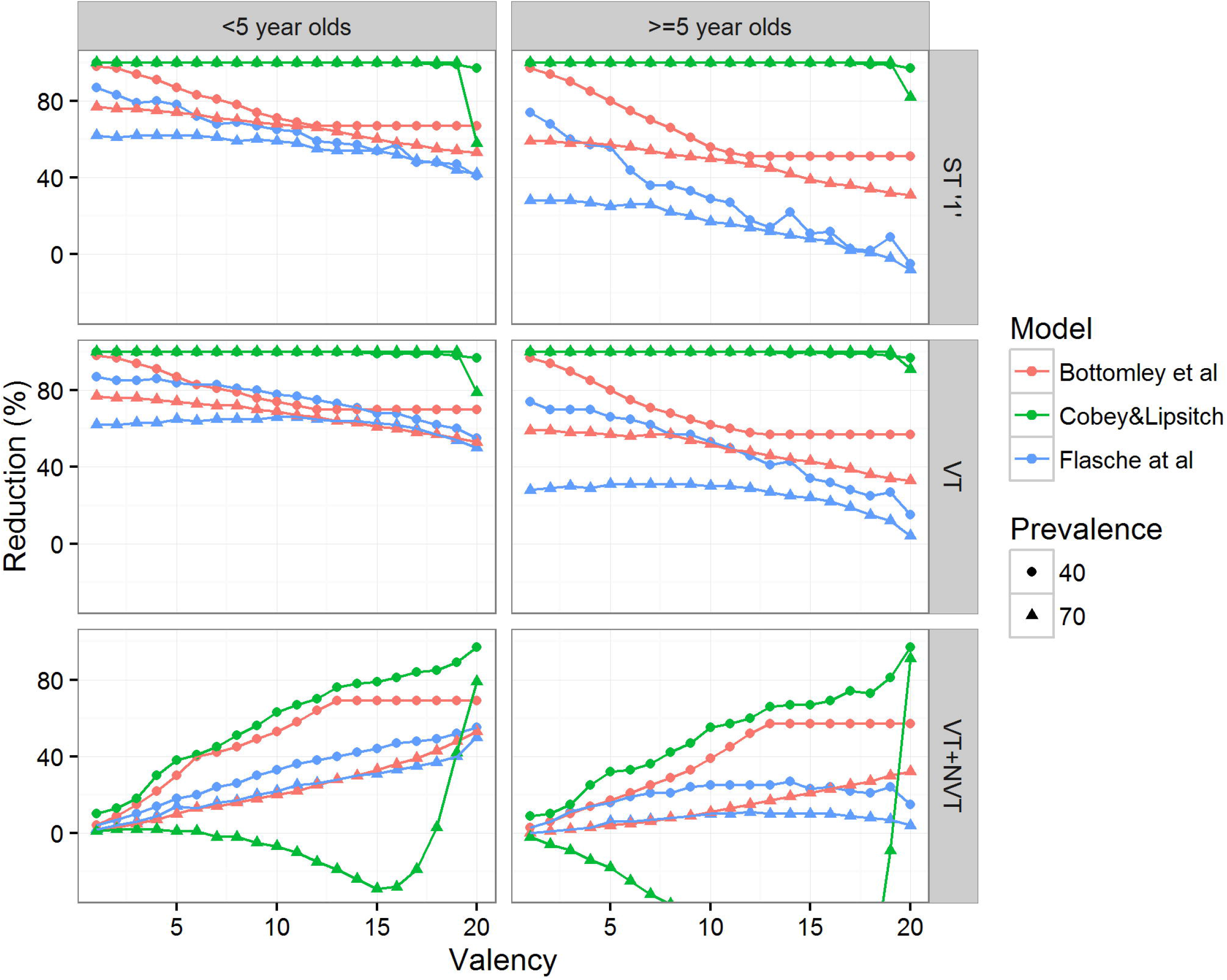
Predicted percentage reduction in the prevalence of pneumococcal carriage (bottom row), VT carriage (middle row) and carriage of the most competitive serotype (top row) 10 years after vaccine introduction, assuming 55% efficacy against acquisition of pneumococcal VTs and 100% coverage.

The impact of vaccination on all serotype carriage measured 10 years after the start of vaccination with a vaccine with 55% effective coverage generally increased with increasing valency. However, for the high transmission scenario M_CL predicted a negative vaccine impact, i.e. an increase in overall pneumococcal carriage through the inclusion of almost all serotypes into the vaccine formulation and only for valencies of 19 and higher predicted a positive impact of vaccination, i.e. a reduction in all serotype carriage prevalence.

The PCV7-like and PCV10-like as well as the PCV13-like and PCV15-like vaccines were indistinguishable because the global prevalence rank of the respectively added serotypes were larger than the number of serotypes considered in this analysis and hence omitted. All three models predict that the impact of vaccination against pediatric VT carriage is similar (<5% difference) across the PCV-like formulation (Figure 3). However, changes in vaccine impact on carriage with any serotype were more pronounced but followed the dynamics of the generic vaccine; i.e. inclusion of more serotypes further reduced carriage prevalence, except in M_CL in the high transmission scenario.

**Fig. 3.**
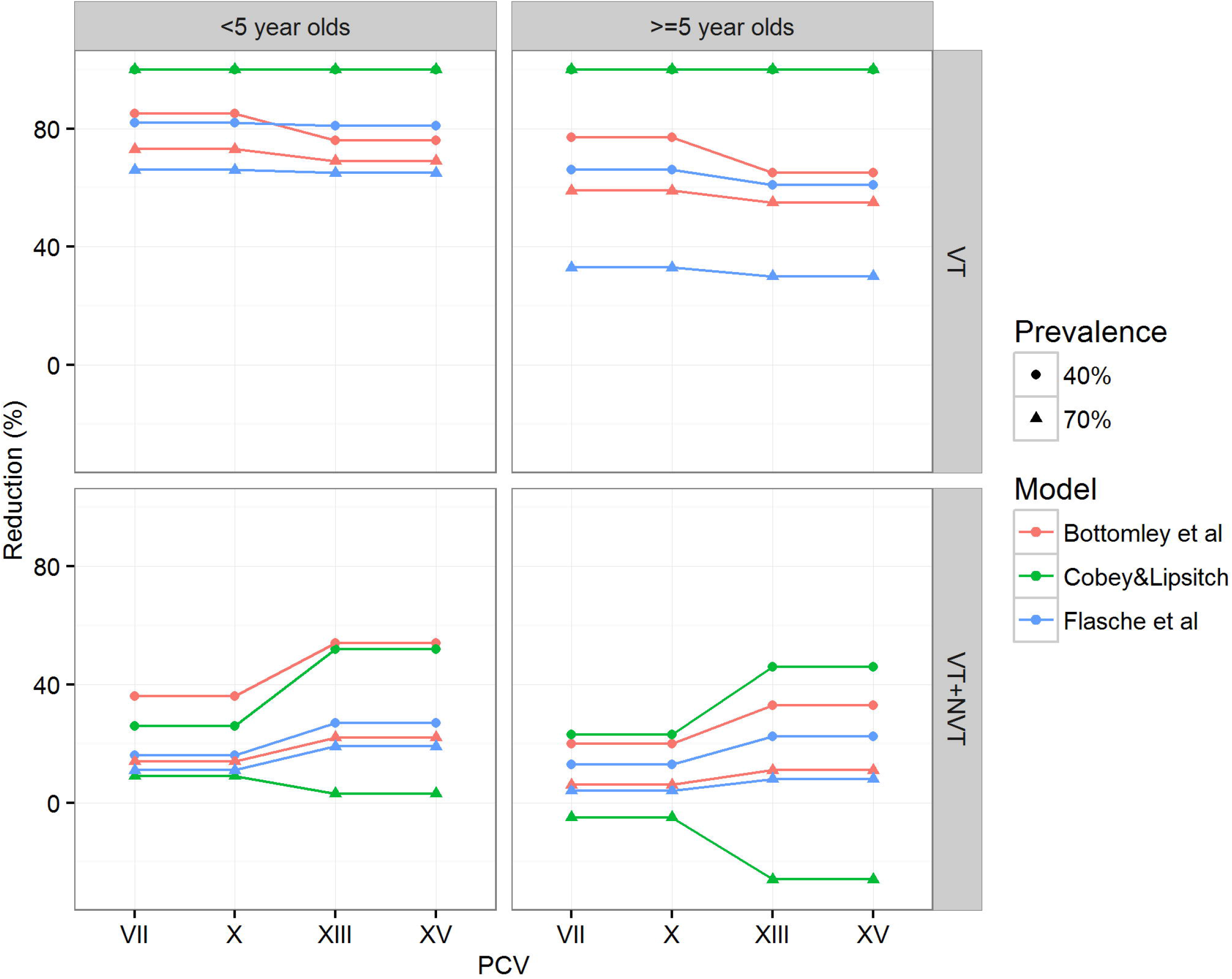
Percentage reduction in the prevalence of pneumococcal carriage 10 years after introduction of a PCV like vaccine, assuming 55% efficacy against acquisition of pneumococcal VTs and 100% coverage.

Consistent with the effect of increasing valency on vaccine impact, increasing the valency of the generic vaccine formulation was predicted to increase the effective coverage needed to eliminate VT carriage in both children and older individuals and in both moderate and high transmission intensity settings (Figure 4). In all three models VT elimination in the general population required less than 10% additional effective coverage to elimination VT carriage among children. In contrast to the other models M_CL predicted that addition of up to half of all serotypes into the generic vaccine formulation would not have a profound effect on the effective coverage needed for elimination of VT carriage but that only inclusion of at least 10 serotypes or 15 serotypes in the moderate and high transmission scenario respectively would. Elimination of pneumococci using a panvalent vaccine was impossible in M_F and also for the high transmission scenario in M_B.

**Fig. 4.**
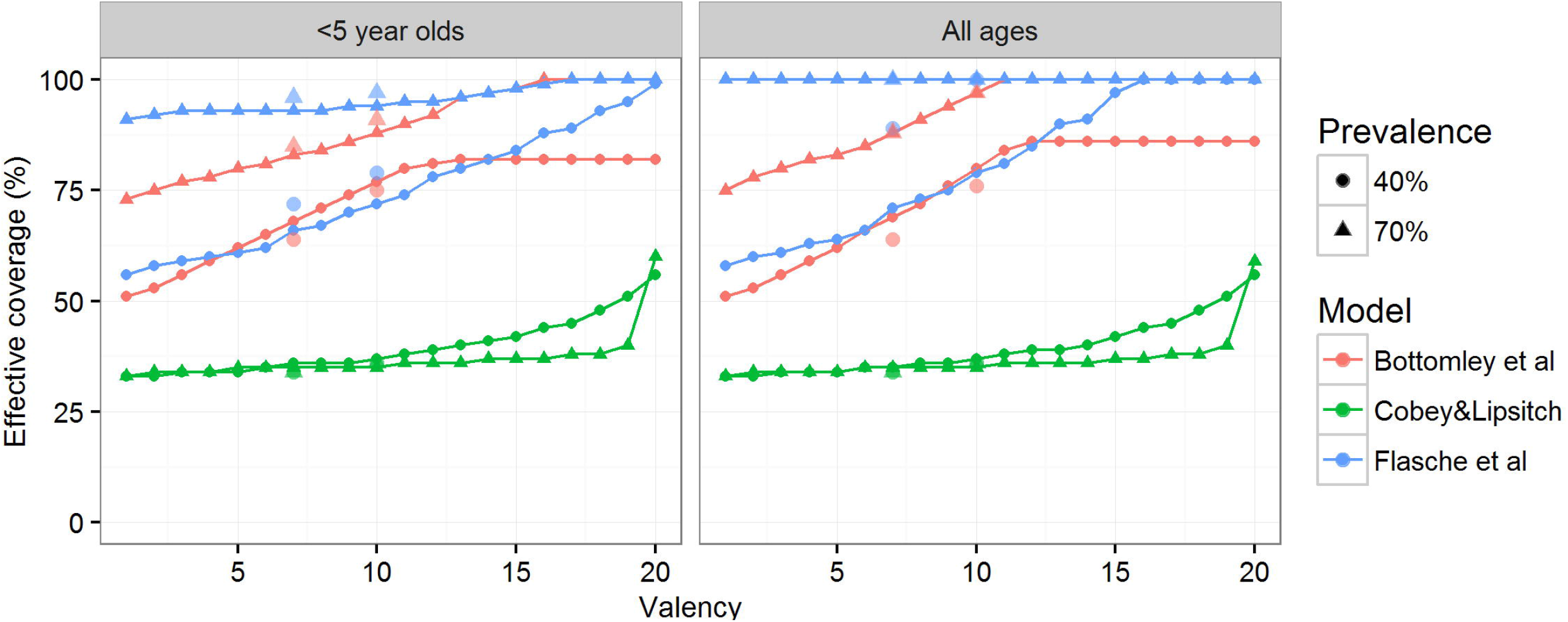
The effective coverage needed to prevent 95% of VT carriage 10 years after the start of vaccination, assuming serotypes are added to the vaccine in order of their carriage prevalence. At valency 7 and 10 the respective effective coverage for PCV7/10 - like and PCV13/15 - like vaccines are indicated by large dots and triangles in respective colors.

For PCV-like vaccines a similar qualitative behavior to the generic vaccine formulations was predicted. The three models predicted that the effective coverage needed to eliminate VT carriage in the population increases by 12, 2 and 11% respectively for the moderate transmission intensity setting if a PCV13 or 15 - like vaccine was used instead of a PCV7 or 10 - like vaccine. In the high transmission intensity setting elimination of VT carriage for the two modelled vaccine formulations was impossible in M_F, required effective coverage larger than 88% and 97% in M_B and required effective coverage of 34% and 37% in M_CL.

## Discussion

Pneumococcal conjugate vaccines have substantially reduced the burden of pneumococcal disease worldwide. However, through replacement with serotypes not targeted by the vaccines a sizeable burden remains and has led to ongoing development of vaccines with higher valency. Using qualitative results across three pneumococcal models that span a variety of assumptions on the acquisition of pneumococcal immunity and serotype competition, we here show that serotype competition from NVTs aids VT elimination. Accordingly, we show that targeting an increasing number of serotypes increases the requirements on vaccine efficacy and/or vaccine coverage to achieve elimination of VT carriage. We predict that the relatively small differences in the number of serotypes targeted by current PCV formulations are unlikely to be substantial enough to lead to measurable differences in the ease of VT elimination but that vaccines that target almost all serotypes may allow continued circulation of the most competitive serotypes even if such vaccine was given at high coverage and the vaccine efficacy against carriage was improved over the efficacy of PCVs.

Given the qualitative nature of this comparison we have only assessed the impact of vaccine valency on the potential of elimination of pneumococcal carriage. The implications of the results for the disease impact of a switch from current PCV formulations to a pan valent vaccine are complex. Increasing vaccine valency could lead to a net increase in pneumococcal disease burden if the highly competitive serotypes are controlled only through a vaccine with limited valency and are also highly pathogenic while the types not targeted by the formulation with limited valency rarely cause disease. In this case, the small disease benefit of controlling more serotypes could be outweighed by the increased circulation of the serotypes in the limited-valency vaccine. The expansion of PCV formulations thus far to incorporate additional serotypes responsible for significant amounts of disease has tended to emphasize highly invasive serotypes, thereby minimizing the potential problem we highlight for expansions of the valency of PCVs. A recent study suggests a method for such that would continue this beneficial approach [29].

In vaccinated persons, these unintended effects might be fully or partially offset through the additional direct vaccine protection against disease, given that PCV formulations thus far have provided >80% protection against disease, with lower efficacy against carriage [28,30–32], This implies that unintended effects of increased valency of vaccines might be of greatest concern - and thus most deserving of surveillance - in age groups within a population that have not been vaccinated, such as healthy adults in most countries at present. We emphasize that the model-comparison exercise here was designed to assess general trends in the behaviors of the models, rather than to predict specifically how a higher-valency vaccine would act in a particular population. Setting-specific model parameterization is required to allow quantification of the differential impact of vaccine of varying valency on pneumococcal disease.

We show that the observation that serotype competition aids VT elimination and hence that increasing the valency of pneumococcal conjugate vaccine is likely to increase the herd immunity threshold is insensitive to different assumptions of peadiatric carriage prevalence and vaccine formulation which we consistently explored across all three models. Most importantly we show that this finding is also insensitive to alternative model assumptions of pneumococcal ecology, the mechanisms of between serotype competition and differences in underlying demographics.

We made the simplifying assumption that serotype specific direct vaccine effects are the same across vaccines and targeted serotypes and that they follow those of current PCV formulations; i.e. an approximate 55% efficacy against carriage acquisition of any targeted serotype [28]. However, because of the complexities involved in the conjugation procedure of PCVs it is unlikely that using current techniques PCVs will be able to target more than 20 of the over 90 pneumococcal serotypes [33]. Vaccines that target common proteins rather than specific capsules on the other hand may prevent pneumococcal disease by different mechanisms, e.g. enhanced IL-17A mediated nasopharyngeal clearance rather than prevention of acquisition [34], While inference of the differential population impact of specific pneumococcal vaccines would require a more precise parameterization including the focus on a specific setting, the qualitative results of this work are likely to similarly apply.

In a few instances the models predicted changes to pneumococcal ecology following vaccination that seem counter-intuitive at first. In the high transmission scenario, M_CL predicted that overall pediatric carriage prevalence would stay relatively constant (less than 5% change) for vaccines that included up to 8 of the most competitive serotypes, but then increase by up to 40% if more serotypes were targeted by the vaccine to finally reduce overall carriage by targeting at least 19 serotypes (Figure 2). This is unique to this model because of its inclusion of a gradient in type-specific ability to prevent additional acquisition: carriers of highly-competitive serotypes are more protected against acquisition of further serotypes. By protecting against the most competitive serotypes through vaccination the remaining serotypes are under less pressure from competition to a point where they act almost independently. For vaccines that target between 9 and 18 of all serotypes in the high transmission setting the prevalence of untargeted serotype in the virtual absence of competition then adds up to exceed the overall prevalence before vaccination (Appendix Figure 1). Furthermore, in M_CL the impact of vaccination with low valency vaccines is similar in both transmission settings but higher for moderate to high valency vaccines in high transmission settings than in moderate transmission settings. In contrast, most models including M_B and M_F predict that transmission intensity and the herd immunity threshold are always positively correlated [35]. In M_CL that same effect is evident only for vaccines that target all serotypes. For vaccines that target most but not all pneumococci this model predicts that the relatively weak competitive pressure from NVTs in combination with a substantial increase in overall pneumococcal prevalence in high transmission scenarios helps to control VT circulation better than vaccination in a moderate transmission scenario with less transmissible VTs but also lower NVT carriage prevalence that compete with VTs. M_F predicted that, assuming 55% effective coverage in the high transmission scenario, increasing the valency to up to 15 serotypes leads to a small increase in vaccine impact on VT carriage which is qualitative different from all other models and scenarios presented here. This is a result of comparing the impact of vaccination on a different number of serotypes. In particular, reduction in carriage prevalence of e.g. serotype “1” as a result of vaccination in the high transmission scenario steadily declines with increasing valency in M_F (Figure 2). However, the indirect effect of vaccination against serotypes of lower prevalence is greater and hence comparing the impact of vaccination on all vaccine serotypes includes both the counter-acting trends. Only in the high transmission setting in M_F is a net increase in vaccine effects against VTs predicted for low valency vaccines.

While the models agree well on the qualitative relation between vaccine valency and the herd immunity threshold we have observed stark differences in the quantitative results. For example M_CL predicted that much lower effective coverage is required for elimination of VT carriage. Many factors contribute to this observation and some could be addressed by more detailed harmonization of the models to a specific setting. However, two intrinsic model assumptions likely drive this behavior: 1) the strength of vaccine protection and 2) the strength of serotype competition. M_CL assumes a relatively strong vaccine protection by reducing susceptibility to VT acquisition permanently after vaccination. M_F assumes vaccine protection, albeit nonleaky, to only hold for 10 years while M_B also assumes lifelong protection from vaccination, although modelled as all-or-nothing and hence even stronger than in M_CL. While the evidence suggest PCVs to be leaky [36] little is known about upcoming pan-valent vaccines. Vaccine protection has been found to remain present 5 years after completion after childhood immunization [37] but there is some evidence that protection declines over time albeit with a half-life that exceeds 5 years [28]. Further, M_CL has the weakest serotype competition of the models by assuming leaky protection against heterologous acquisition of additional colonizing strains during carriage with the most competitive serotypes providing stronger protection against new acquisition. In comparison M_B and M_F respectively assume competitive exclusion on acquisition and potential co-infection but only after non-leaky heterologous immunity following acquisition has ended. With the relatively weak competition of serotypes in M_CL a targeted pediatric carriage prevalence is achieved with lower transmission intensity and this in turn will ease elimination of vaccine serotypes in comparison to the other models where transmission is more intense. Recent advances in molecular serotyping methods have shown the pneumococci frequently co-colonise [38,39] and while epidemiological studies suggest that carriage induces both homologous and heterologous protection such protection from a single episode of carriage likely is relatively weak [20,40,41],

## Conclusion

Using three different modelling approaches for pneumococcal ecology that represent a range of alternative assumptions on pneumococcal immunity and serotype competition we found that NVT competition helps vaccines of limited valencies eliminate VT carriage. This implies that new vaccines that targeted the majority of pneumococcal serotypes will benefit less from NVT competition and are likely to offer less indirect protection than current PCVs. Head to head comparison of current PCVs with high-valency vaccines should not only be on the grounds of non-inferiority of direct effects but should also account for indirect effects, and closely monitor IPD endpoints.

## Appendix

**Appendix Figure 1:**
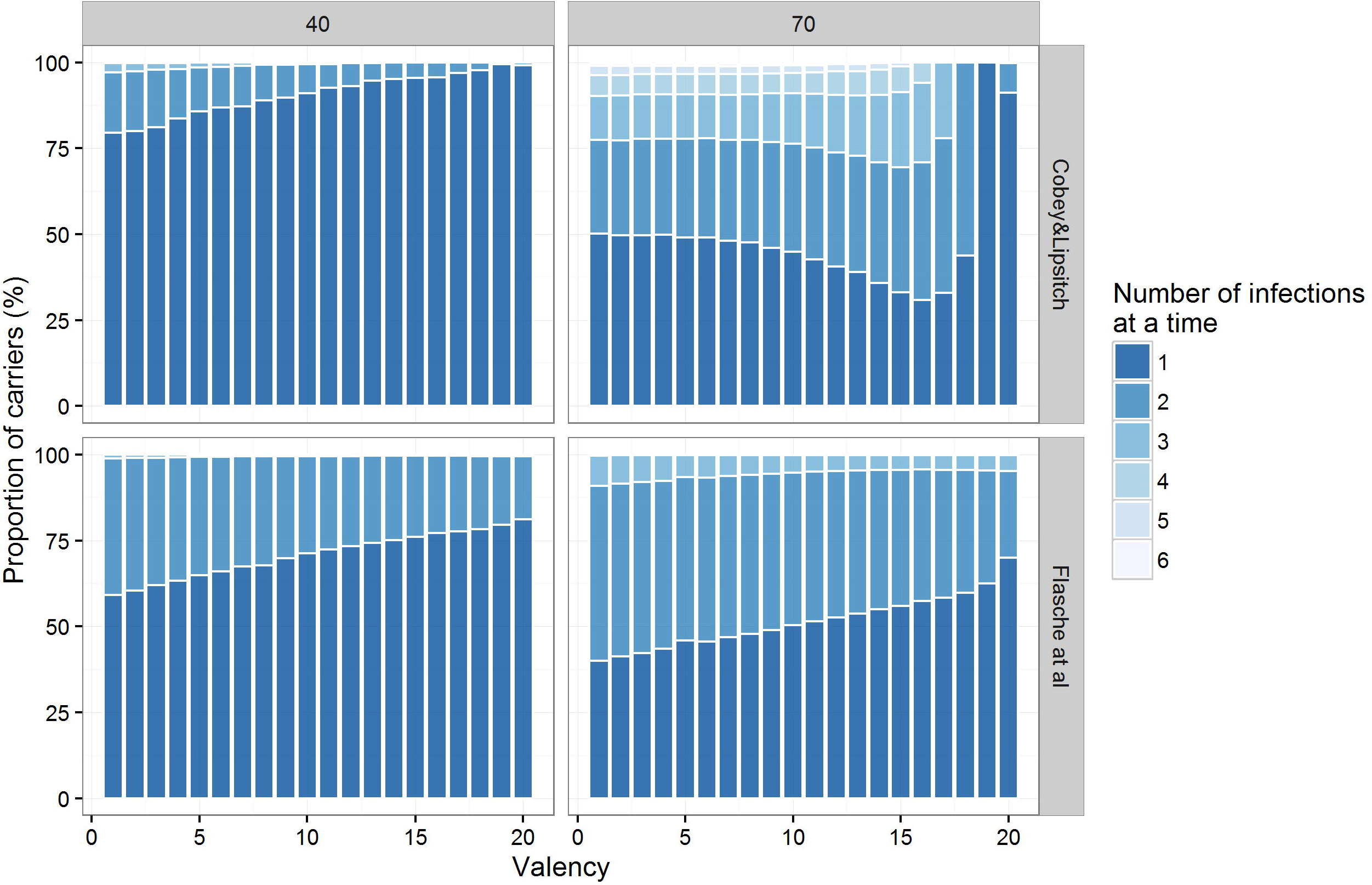
The proportion of carriers that carry 1,2…,6 pneumococci at a time following ten years of vaccination. Rows show different models and columns different transmission intensities indicated by the prevalence of carriage in less than 5-year-old children before vaccination. In M_B each host can only carry one pneumococci at a time and hence the model is not represented here.

## References

1. Centers for Disease Control and Prevention (CDC) 2016 Pink book - Pneumococcal disease.

2. Waight, P. A., Andrews, N. J., Ladhani, S. N., Sheppard, C. L., Slack, M. P. E. & Miller, E. 2015 Effect of the 13-valent pneumococcal conjugate vaccine on invasive pneumococcal disease in England and Wales 4 years after its introduction: an observational cohort study. Lancet Infect. Dis. 3099, 1–9. (doi:10.1016/S1473-3099(15)70044-7)

3. Moore, M. R. et al. 2016 Effectiveness of 13-valent pneumococcal conjugate vaccine for prevention of invasive pneumococcal disease in children in the USA: a matched case-control study. Lancet Respir. Med. 4, 399–406. (doi:10.1016/S2213-2600(16)00052-7)

4. Angoulvant, F. et al. 2014 Early Impact of 13-Valent Pneumococcal Conjugate Vaccine on Community-Acquired Pneumonia in Children. Clin. Infect. Dis., 1–7. (doi:10.1093/cid/ciu006)

5. Harboe, Z. & Dalby, T. 2014 Impact of 13-Valent Pneumococcal Conjugate Vaccination in Invasive Pneumococcal Disease Incidence and Mortality. … Infect. Dis. 59,1066–1073. (doi:10.1093/cid/ciu524)

6. Rückinger, S., van der Linden, M., Siedler, A. & von Kries, R. 2011 Potential Benefits From Currently Available Three Pneumococcal Vaccines for Children - Population-Based Evaluation. Klin. Padiatr. (doi:10.1055/s-0030-1268465)

7. Weiss, S., Falkenhorst, G., van der Linden, M., Imohl, M. & von Kries, R. 2015 Impact of 10- and 13- valent pneumococcal conjugate vaccines on incidence of invasive pneumococcal disease in children aged under 16 years in Germany, 2009 to 2012. Euro Surveill. Bull. Eur. sur les Mal. Transm. = Eur. Commun. Dis. Bull. 20, 21057.

8. Olarte, L. et al. 2015 Impact of the 13-Valent Pneumococcal Conjugate Vaccine on Pneumococcal Meningitis in U.S. Children. Clin. Infect. Dis. (doi:10.1093/cid/civ368)

9. Greenberg, D., Givon-Lavi, N., Ben-Shimol, S., Ziv, J. B. & Dagan, R. 2015 Impact of PCV7/PCV13 introduction on community-acquired alveolar pneumonia in children <5 years. Vaccine 33, 4623–4629. (doi:10.1016/j.vaccine.2015.06.062)

10. Bruce, M. G., Singleton, R., Bulkow, L., Rudolph, K., Zulz, T., Gounder, P., Hurlburt, D., Bruden, D. & Hennessy, T. 2015 Impact of the 13-valent pneumococcal conjugate vaccine (pcv13) on invasive pneumococcal disease and carriage in Alaska. Vaccine 33, 4813–4819. (doi:10.1016/j.vaccine.2015.07.080)

11. Ben-Shimol, S., Givon-Lavi, N., Leibovitz, E., Raiz, S., Greenberg, D. & Dagan, R. 2016 Impact of Widespread Introduction of Pneumococcal Conjugate Vaccines on Pneumococcal and Nonpneumococcal Otitis Media. Clin. Infect. Dis., ciw347. (doi:10.1093/cid/ciw347)

12. Feikin, D. R. et al. 2013 Serotype-specific changes in invasive pneumococcal disease after pneumococcal conjugate vaccine introduction: a pooled analysis of multiple surveillance sites. PLoS Med. 10, el001517. (doi:10.1371/journal.pmed. 1001517)

13. Weinberger, D. M., Malley, R. & Lipsitch, M. 2011 Serotype replacement in disease after pneumococcal vaccination. Lancet 6736, 1–12. (doi:10.1016/S0140-6736(10)62225-8)

14. Alderson, M. R. 2016 Status of vaccine research and development of pediatric vaccines for Streptococcus pneumoniae. Vaccine (doi:10.1016/j.vaccine.2016.03.107)

15. Flasche, S., Edmunds, W. J., Miller, E., Goldblatt, D., Robertson, C. & Choi, Y. H. 2013 The impact of specific and non-specific immunity on the ecology of Streptococcus pneumoniae and the implications for vaccination. Proc. R. Soc. B Biol. Sci. 280, 20131939–20131939. (doi:10.1098/rspb.2013.1939)

16. Bottomley, C., Roca, A., Hill, P. C., Greenwood, B. & Isham, V. 2013 A mathematical model of serotype replacement in pneumococcal carriage following vaccination. J. R. Soc. Interface 10, 20130786. (doi:10.1098/rsif.2013.0786)

17. Lipsitch, M. 1997 Vaccination against colonizing bacteria with multiple serotypes. Proc. Natl. Acad. Sci. U. S. A. 94, 6571–6.

18. Trzciński, K., Li, Y., Weinberger, D. M., Thompson, C. M., Cordy, D., Bessolo, A., Malley, R. & Lipsitch, M. 2015 Effect of Serotype on Pneumococcal Competition in a Mouse Colonization Model. MBio 6, e00902–15. (doi:10.1128/mBio.00902-15)

19. Paterson, G. K. & Mitchell, T. J. 2006 Innate immunity and the pneumococcus. Microbiology 152, 285–293. (doi:10.1099/mic.0.28551-0)

20. Granat, S. M., Ollgren, J., Herva, E., Mia, Z., Auranen, K. & Mäkelä, P. H. 2009 Epidemiological evidence for serotype-independent acquired immunity to pneumococcal carriage. J. Infect. Dis. 200, 99–106. (doi:10.1086/599364)

21. Malley, R., Trzcinski, K., Srivastava, A., Thompson, C. M., Anderson, P. W. & Lipsitch, M. 2005 CD4+ T cells mediate antibody-independent acquired immunity to pneumococcal colonization. Proc. Natl. Acad. Sci. 102, 4848–4853. (doi:10.1073/pnas.0501254102)

22. Lu, Y.-J. et al. 2008 Interleukin-17A mediates acquired immunity to pneumococcal colonization. PLoS Pathog. 4, e1000159. (doi:10.1371/journal.ppat.1000159)

23. Cobey, S. & Lipsitch, M. 2012 aNiche and Neutral Effects of Acquired Immunity Permit Coexistence of Pneumococcal Serotypes. Science (80-.). 335, 1376–1380. (doi:10.1126/science.1215947)

24. Choi, Y. H., Jit, M., Gay, N., Andrews, N., Waight, P. a, Melegaro, A., George, R. & Miller, E. 2011 7-Valent pneumococcal conjugate vaccination in England and Wales: is it still beneficial despite high levels of serotype replacement? PLoS One 6, e26190. (doi:10.1371/journal.pone.0026190)

25. Choi, Y. H. Y. H., Jit, M., Flasche, S., Gay, N. & Miller, E. 2012 Mathematical modelling long-term effects of replacing Prevnar7 with Prevnar13 on invasive pneumococcal diseases in England and Wales. PLoS One 7, e39927. (doi:10.1371/journal.pone.0039927)

26. Hanage, W. P., Finkelstein, J. A., Huang, S. S., Pelton, S. I., Stevenson, A. E., Kleinman, K., Hinrichsen, V. L. & Fraser, C. 2010 Evidence that pneumococcal serotype replacement in Massachusetts following conjugate vaccination is now complete. Epidemics 2, 80–84. (doi:10.1016/j.epidem.2010.03.005)

27. Le Polain de Waroux, O., Edmunds, W. J. & Flasche, S. 2016 RESPICAR. Prep.

28. Le Polain De Waroux, O., Flasche, S., Prieto-Merino, D., Goldblatt, D. & Edmunds, W. J. 2015 The Efficacy and Duration of Protection of Pneumococcal Conjugate Vaccines Against Nasopharyngeal Carriage. Pediatr. Infect. Dis.J. 34, 858–864. (doi:10.1097/INF.0000000000000717)

29. Nurhonen, M. & Auranen, K. 2014 Optimal Serotype Compositions for Pneumococcal Conjugate Vaccination under Serotype Replacement. PLoS Comput. Biol. 10, el003477. (doi:10.1371/journal.pcbi. 1003477)

30. Tregnaghi, M. W. et al. 2014 Efficacy of pneumococcal nontypable Haemophilus influenzae protein D conjugate vaccine (PHiD-CV) in young Latin American children: A double-blind randomized controlled trial. PLoS Med. 11, e1001657. (doi:10.1371/journal.pmed. 1001657)

31. Cutts, F. T. et al. 2005 Efficacy of nine-valent pneumococcal conjugate vaccine against pneumonia and invasive pneumococcal disease in The Gambia: randomised, double-blind, placebo-controlled trial. Lancet 365, 1139–46. (doi:10.1016/S0140-6736(05)71876-6)

32. O’Brien, K. L. et al. 2003 Efficacy and safety of seven-valent conjugate pneumococcal vaccine in American Indian children: group randomised trial. Lancet 362, 355–61. (doi:10.1016/S0140- 6736(03)14022-6)

33. Rodgers, G. L. & Klugman, K. P. 2011 The future of pneumococcal disease prevention. Vaccine 29 Suppl 3, C43–8. (doi:10.1016/j.vaccine.2011.07.047)

34. Malley, R. & Anderson, P. W. 2012 Serotype-independent pneumococcal experimental vaccines that induce cellular as well as humoral immunity. Proc. Natl. Acad. Sci. U. S. A. 109, 1–5. (doi:10.1073/pnas.1121383109)

35. Fine, P., Eames, K. & Heymann, D. L. 2011 ‘Herd Immunity’: A Rough Guide. Clin. Infect. Dis. 52, 911916. (doi:10.1093/cid/cir007)

36. Dagan, R., Givon-Lavi, N., Fraser, D., Lipsitch, M., Siber, G. R. & Kohberger, R. 2005 Serum serotype-specific pneumococcal anticapsular immunoglobulin g concentrations after immunization with a 9- valent conjugate pneumococcal vaccine correlate with nasopharyngeal acquisition of pneumococcus. J. Infect. Dis. 192, 367–376. (doi:10.1086/431679)

37. Madhi, S. a, Adrian, P., Kuwanda, L., Cutland, C., Albrich, W. C. & Klugman, K. P. 2007 Long-term effect of pneumococcal conjugate vaccine on nasopharyngeal colonization by Streptococcus pneumoniae--and associated interactions with Staphylococcus aureus and Haemophilus influenzae colonization--in HIV-Infected and HIV-uninfected children. J. Infect. Dis. 196, 1662–6. (doi:10.1086/522164)

38. Turner, P., Hinds, J., Turner, C., Jankhot, A., Gould, K., Bentley, S. D., Nosten, F. & Goldblatt, D. 2011 Improved detection of nasopharyngeal co-colonization by multiple pneumococcal serotypes using latex agglutination or molecular serotyping by microarray. J. Clin. Microbiol. 49, 1784–1789. (doi:10.1128/JCM.00157-11)

39. Satzke, C., Dunne, E. M., Porter, B. D., Klugman, K. P. & Mulholland, E. K. 2015 The PneuCarriage Project: A Multi-Centre Comparative Study to Identify the Best Serotyping Methods for Examining Pneumococcal Carriage in Vaccine Evaluation Studies. PLOS Med. 12, e1001903. (doi:10.1371/journal.pmed.1001903)

40. Weinberger, D. M., Dagan, R., Givon-Lavi, N., Regev-Yochay, G., Malley, R. & Lipsitch, M. 2008 Epidemiologic evidence for serotype-specific acquired immunity to pneumococcal carriage. J. Infect. Dis. 197, 1511–8. (doi:10.1086/587941)

41. Lipsitch, M., Whitney, C. G., Zell, E., Kaijalainen, T., Dagan, R. & Malley, R. 2005 Are anticapsular antibodies the primary mechanism of protection against invasive pneumococcal disease? PLoS Med. 2, el5. (doi:10.1371/journal.pmed.0020015)

42. Lipsitch, M., Colijn, C., Cohen, T., Hanage, W. P. & Fraser, C. 2009 No coexistence for free: neutral null models for multistrain pathogens. Epidemics 1, 2. (doi:10.1016/j.epidem.2008.07.001)

